# Multi-omics data and model integration reveal the main mechanisms associated with respiro-fermentative metabolism and ethanol stress responses in *Kluyveromyces marxianus*

**DOI:** 10.1101/2024.06.06.597719

**Authors:** Maurício Alexander de Moura Ferreira, Wendel Batista da Silveira

## Abstract

*Kluyveromyces marxianus* is a yeast capable of fermenting sugars into ethanol and growing at high temperatures (>37ºC). However, it is less tolerant to ethanol than *Saccharomyces cerevisiae*, which limits its application in second-generation ethanol production. Since the mechanisms of ethanol stress response are still poorly described, especially compared to *S. cerevisiae*, we used an integrative multi-omics approach, combining transcriptomics, coexpression networks, gene regulation, and genome-scale metabolic modelling to gain insights about these mechanisms. Through metabolic modelling, we predicted the occurrence of a respiro-fermentative metabolism and its onset as the dilution rate increased. From gene coexpression networks, we detected that the protein quality control system is a main mechanism involved in the ethanol stress response. Further, we identified key regulators in the ethanol stress response, such as *HAP3, MET4*, and *SNF2*, and assessed how disturbances in their gene expression affect cellular metabolism. We also found that amino acid metabolism, membrane lipid metabolism, and ergosterol exhibit increased metabolic flux under the explored conditions. These findings provide useful cues to develop and implement genetic and metabolic engineering strategies to enhance ethanol tolerance.

## 1 Introduction

Climate change has driven the development of new biofuel production processes to mitigate the damage caused by greenhouse gas emissions from burning fossil fuels [1–3]. Ethanol, the main biofuel marketed globally [4], is primarily produced from sugarcane and starch-based raw materials (first-generation ethanol) [3]. However, this process competes with food production. In this context, producing ethanol from alternative sources has been considered essential to avoid using arable land, water resources, and fertilisers, and to prevent competition with food production [5]. These alternative sources are generally industrial by-products, such as whey, which is rich in lactose, and lignocellulosic biomass, which is rich in glucose, xylose, mannose, and arabinose.

First-generation ethanol is produced by the yeast *Saccharomyces cerevisiae* [6] due to its remarkable ability to produce high ethanol titres from glucose and sucrose. At high sugar concentrations, enzymes of the central metabolism have their abundance altered, favouring enzymes that catalyse fermentative metabolism over respiratory metabolism [7]. This phenomenon, known as overflow metabolism (or the Crabtree effect in yeasts), allows *S. cerevisiae* to produce ethanol by fermentation even under aerobic conditions [8]. Despite its performance, *S. cerevisiae* is unable to assimilate sugars found in abundant feedstocks such as lactose and xylose [5]. Moreover, *S. cerevisiae* does not tolerate high temperatures, making difficult its application for second-generation ethanol production thorough processes of simultaneous saccharification and fermentation of lignocellulosic biomasses, which generally occur between 50 and 60ºC [9]. On the other hand, *Kluyveromyces marxianus* is a thermotolerant yeast capable of fermenting sugars at temperatures ranging from 42 to 48ºC [10]. Additionally, it can use lactose and xylose as carbon and energy sources. Although it can produce ethanol from sugars such as lactose and glucose, it is not considered a Crabtree-positive yeast [11]. In addition, *K. marxianus* is less tolerant to high ethanol concentrations, which is a drawback for its application in ethanol production [12,13]. Thus, efforts have been made to uncover the mechanisms of ethanol stress response in this yeast and to obtain strains with improved tolerance [14].

Studies on ethanol stress in yeasts have primarily focused on *S. cerevisiae*. There are still few studies on non-*Saccharomyces* yeasts. In *K. marxianus*, physiological responses to stress conditions are largely inferred from results with its sister species, *K. lactis*, and with *S. cerevisiae* [14]. However, there are fundamental differences between the species, highlighting the need for a better understanding of stress response mechanisms in *K. marxianus*.

Specifically, during ethanol stress, *S. cerevisiae* induces an increase in ergosterol and unsaturated fatty acids in the plasma membrane, while in *K. marxianus*, this response relies on the strain and cultivation condition [15]. Regarding gene expression, different sets of genes are induced and repressed between the two species. Differences in metabolic profiles between *K. marxianus* and *S. cerevisiae* are also notable [16]. Additionally, there are few large-scale studies at the systems level that integrate both molecular and metabolic knowledge.

Metabolism can be studied computationally through flux balance analysis (FBA) where genome-scale metabolic models (GEMs) are used to predict metabolic flux [17]. GEMs include the stoichiometry of each reaction in the network and information about the genes responsible for the enzymes that catalyse each reaction, allowing a direct association between genotype and phenotype. These models are useful for phenotype simulations and can be improved with additional data, especially catalytic efficiency and enzyme concentration [18,19], and gene expression [20]. GEMs integrated with enzyme constraints (ecGEMs) enable the study of proteome resource allocation by considering enzyme concentration and allow the study of more complex phenotypes, such as metabolic changes that occur at high growth rates and carbon source concentrations. Specifically, ecGEMs allow studying and predicting the Crabtree effect, which occurs in a manner dependent on the allocation of enzymatic resources. In *K. marxianus*, which exhibits respiro-fermentative metabolism [10], there are currently no modelling studies on this respiro-fermentative metabolism. Despite the utility of GEMs and ecGEMs, metabolism is highly dependent on gene expression and regulation, which can be studied through gene co-expression networks (GCNs) and gene regulatory networks (GRNs). These networks can be inferred from transcriptomics data, associating expression patterns of individual genes that occur jointly (in the case of GCNs) [21], or associating the expression of transcription factor (TF) genes with possible target genes through unsupervised or reference-guided clustering (in the case of GRNs) [22].

To address this knowledge gap in *K. marxianus*, here we focus on reconstructing and analysing the enzyme-constrained metabolic network of *K. marxianus*, and characterising GCNs and GRNs using available transcriptomics data. Further, we integrate these different modelling approaches to investigate gene co-expression, gene regulation, metabolism regulation, and resource allocation in response to ethanol stress. Finally, we were able to predict the respiro-fermentative metabolism *K. marxianus* and establish a basis for proposing metabolic engineering strategies to improve tolerance to high ethanol concentrations in *K. marxianus*.

## 2 Material and methods

### 2.1 Integrating enzyme constraints in the *K. marxianus* GEM

To enhance the predictive capacity of the GEM for *K. marxianus*, we integrated catalytic efficiency values (*k*_*cat*_) and enzyme concentrations into the consensus model iSM996 [23]. For this, we used the GECKO Toolbox 3 [24]. Information about the enzymes in the model, such as molecular mass, EC number, and amino acid sequence, was retrieved from UniProt [25] and KEGG [26]. The *k*_*cat*_ values were obtained in two ways: using values included in the BRENDA database [27] and predicted by the DLKcat tool [28]. For each *k*_*cat*_ value of an enzyme obtained from BRENDA or DLKcat, the highest available value was used. To make predictions using DLKcat, the SMILES code of each metabolite was obtained from the PubChem database [29]. Lastly, we allowed flexibilization of the *k*_*cat*_ values for the model to achieve the maximum growth rate of 0.56 h^-1^ using glucose as carbon source. To evaluate the predictive capabilities of the model, we simulated batch cultures using different carbon sources (glucose, fructose, sucrose, galactose, lactose, and xylose) and compared the predictions to experimental data [30–36], with the following optimization set-up:

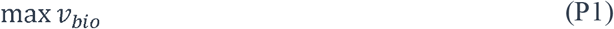

subject to

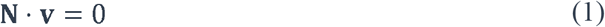

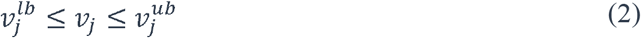

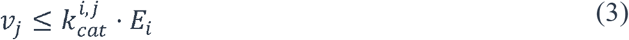

where *v*_*bio*_ is the flux through the biomass pseudoreaction, **N** is the stoichiometric matrix, **v** is the flux distribution vector, 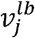 is the flux lower bound of reaction *j, v*_*j*_ is the flux through reaction *j*, 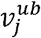is the flux upper bound of reaction *j*, 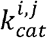 is the *k*_*cat*_ of enzyme *i* catalyzing reaction *j*, and *E* is the concentration of enzyme *i*.

Next, we simulated glucose-limited chemostats with increasing dilution and substrate uptake rates to predict metabolic shifts. We constrained *v*_*bio*_ with respect to increasing values of 0 to 0.5, while minimizing the flux through the carbon source exchange reaction:

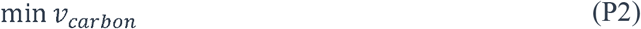

subject to

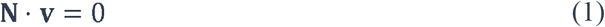

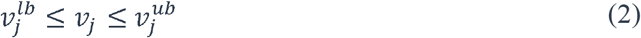

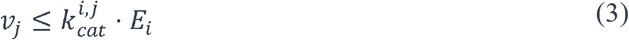

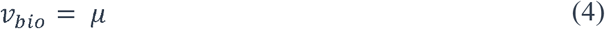

where *v*_*car bon*_ is the flux through the carbon source exchange reaction, and μ is the growth rate. We named the finalized model as eciSM996. All model refinements were performed using the COBRA Toolbox 3 [37] and/or the RAVEN Toolbox 2 [38] in MATLAB (The MathWorks Inc., Natick, Massachusetts).

### 2.2 Obtaining and processing RNAseq data

To investigate the effects of ethanol stress in *K. marxianus*, we used transcriptomic data from Diniz et al. [39] and Mo et al. [40], which were obtained from cultures of *K. marxianus* exposed to ethanol. The data obtained from Diniz et al. [39] were unprocessed FASTQ reads, while Mo et al. [40] provided the gene count matrix. To process the data from Diniz et al. [39], we aligned the reads to the *K. marxianus* reference genome (strain DMKU3-1042) using Bowtie2 [41]. Using these alignments, we generated the count matrix using featureCounts [42]. Lastly, we normalised the counts with DESeq2 [43], using the geometric mean for each gene across all samples, for both the counts generated from Diniz et al. [39] and the matrix obtained from Mo et al. [40].

### 2.3 Inferring gene co-expression and regulation networks

To identify key genes in the ethanol stress response, we first inferred gene co-expression networks (GCNs). For this, we used the BioNERO package [44], available in R/Bioconductor [45]. We used the normalised count matrices as input data and filtered out genes with no expression value and outlier genes. We also removed genes that function as confounding variables. The inferred GCNs were of the hybrid signed type, so only positive associations in the adjacency matrix are considered:

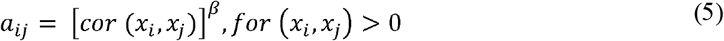

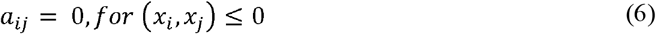

where *a* is the *i*-th, *j*-th element of the adjacency matrix, *x* is the expression value of the gene, and *β* is soft power threshold. Using the identified modules, we annotated them using Gene Ontology (GO) [46] and InterPro functional domains [47]. We selected the inferred modules with the highest correlation with the analysed phenotypes to identify the key genes.

We reconstructed the gene regulatory network (GRN) of *K. marxianus* by concatenating two GRNs. First, we inferred a GRN using BioNERO, with the normalised count matrices as input, identifying specific regulator-target pairs for each input matrix. Then, we generated a second GRN from the *S. cerevisiae* GRN available in the Yeastract database [48]. For each regulator and its respective target genes, we identified the corresponding orthologous genes in *K. marxianus*, substituting the existing gene families in both species and using bidirectional BLAST for genes without a direct association.

### 2.4 Integration of GRNs and metabolic models

To integrate the inferred GRNs with metabolism, we used the Probabilistic Regulation of Metabolism (PROM) algorithm [49]. PROM calculates the probability of a gene being regulated in the absence of its regulator. The relationship between regulator and target is binarized to represent “on” or “off” states for a given target. This probability is represented as:

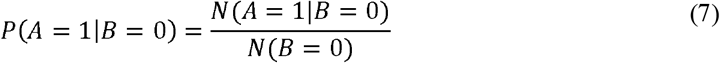

where *P* is the probability, *A* is the target gene, *B* is the regulator, and *N* is the number of occurrences. We used PROM to constrain the conventional iSM996 model by solving the following problem:

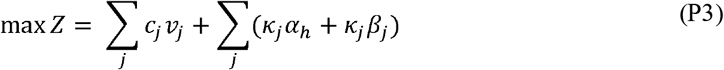

subject to

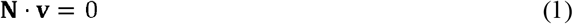

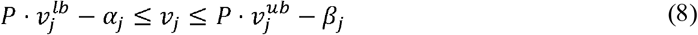

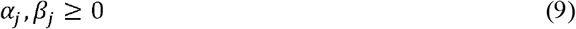

where *c* is the objective coefficient for reaction *j, a* and *β* are adjustments of the lower bound 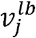 and upper bound 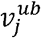, and *P* is the probability calculated in Equation 3. We performed all simulations using the COBRA Toolbox 3 [37] and/or the RAVEN Toolbox 2 [38] in MATLAB (The MathWorks Inc., Natick, Massachusetts).

### 2.5 Integrating gene expression in the *K. marxianus* GEM

To investigate how variations in gene expression affect metabolism, we integrated gene expression data from the count matrices with the metabolic model iSM996 using RIPTiDe [50]. This tool minimises fluxes weighted by the expression of genes controlling the reactions:

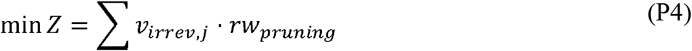

subject to

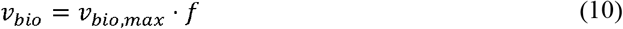

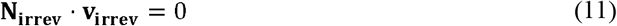

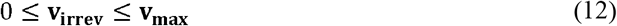

where *v*_*irrev,j*_ is the metabolic flux through an irreversible reaction *j, rw*_*pruning*_ is the weight determined by gene expression, f is the correction factor, **N**_**irrev**_ is the stoichiometric matrix in irreversible format, and **V**_**irrev**_ is the flux vector for irreversible reactions. By obtaining condition-specific models, we maximised biomass production through the reduced network:

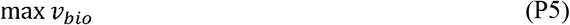

using constraints from Equations 1 and 2.

Next, we applied the predicted flux distribution as constraints in the eciSM996 model to predict resource allocation under these conditions. With this model, we solved problem P5, adding constraint Equation 3 and the following constraint:

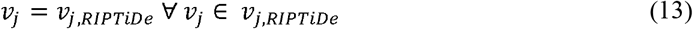

where *v*_*j,RIPTiDe*_ is the previously predicted flux through reaction *j*.

## 3 Results and discussion

### 3.1 The ecGEM eciSM966 predicts respiro-fermentative metabolism

Conventional metabolic models are useful for predicting various phenotypes and physiological conditions, but they cannot simulate metabolic changes such as overflow metabolism or diauxic growth without additional constraints [19,51]. These phenotypes can be predicted by integrating enzymatic constraints into these models, such as *k*_*cat*_ values and enzyme concentrations. In this regard, we used GECKO3 to reconstruct the ecGEM of *K. marxianus* from the conventional GEM iSM966, resulting in eciSM966. This new model has 10,597 reactions, 2,523 metabolites, and 997 genes. Out of the total reactions, 9,293 of them have integrated *k*_*cat*_ values, distributed among 991 enzymes.

Using this model, we simulated batch cultures using different carbon sources. We compared the predicted growth rates to experimental growth rates and those predicted by a conventional GEM (Figure 1). We observed that the eciSM966 model can predict growth rates closer to experimental values than the conventional GEM. Then, we simulated chemostats at different dilution rates to assess metabolic changes associated with high growth rates. We observed that glucose and oxygen uptake, and CO_2_ production, linearly increase along dilution rates up to a rate of 0.35 h^-1^ (Figure 2). From this rate onwards, there is a spike in ethanol production and a decrease in oxygen uptake. However, oxygen continues to be taken up, unlike what occurs in the ecYeastGEM model of *S. cerevisiae* [19]. Additionally, the fraction of total available proteins used also increased up to a dilution rate of 0.35 h^-1^, with a slight decrease after this rate, indicating less enzyme allocation at higher dilution rates.

**Figure 1.**
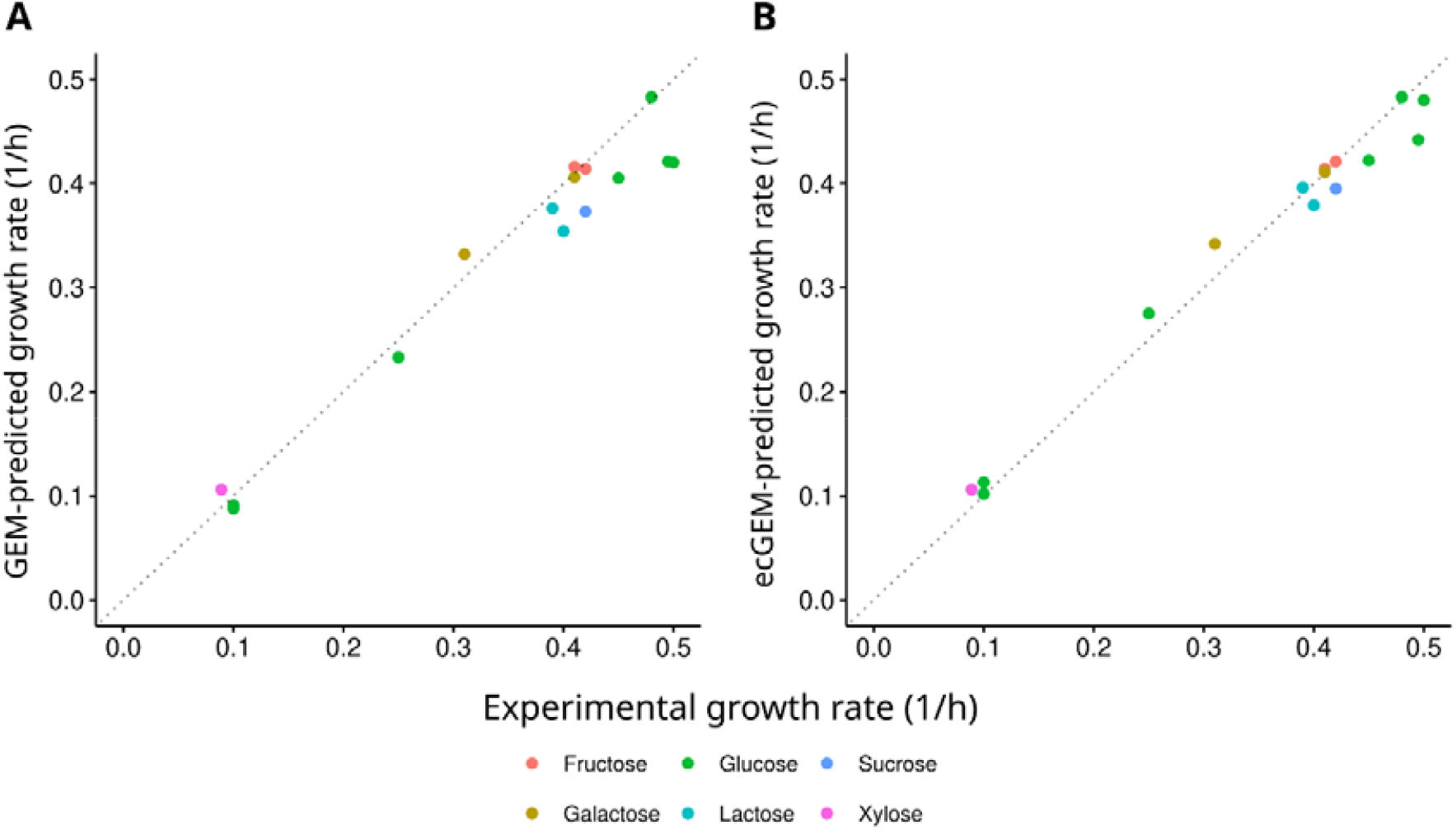
Growth rate comparison between experimental values and predicted values. (A) GEM predictions. (B) ecGEM predictions.

**Figure 2.**
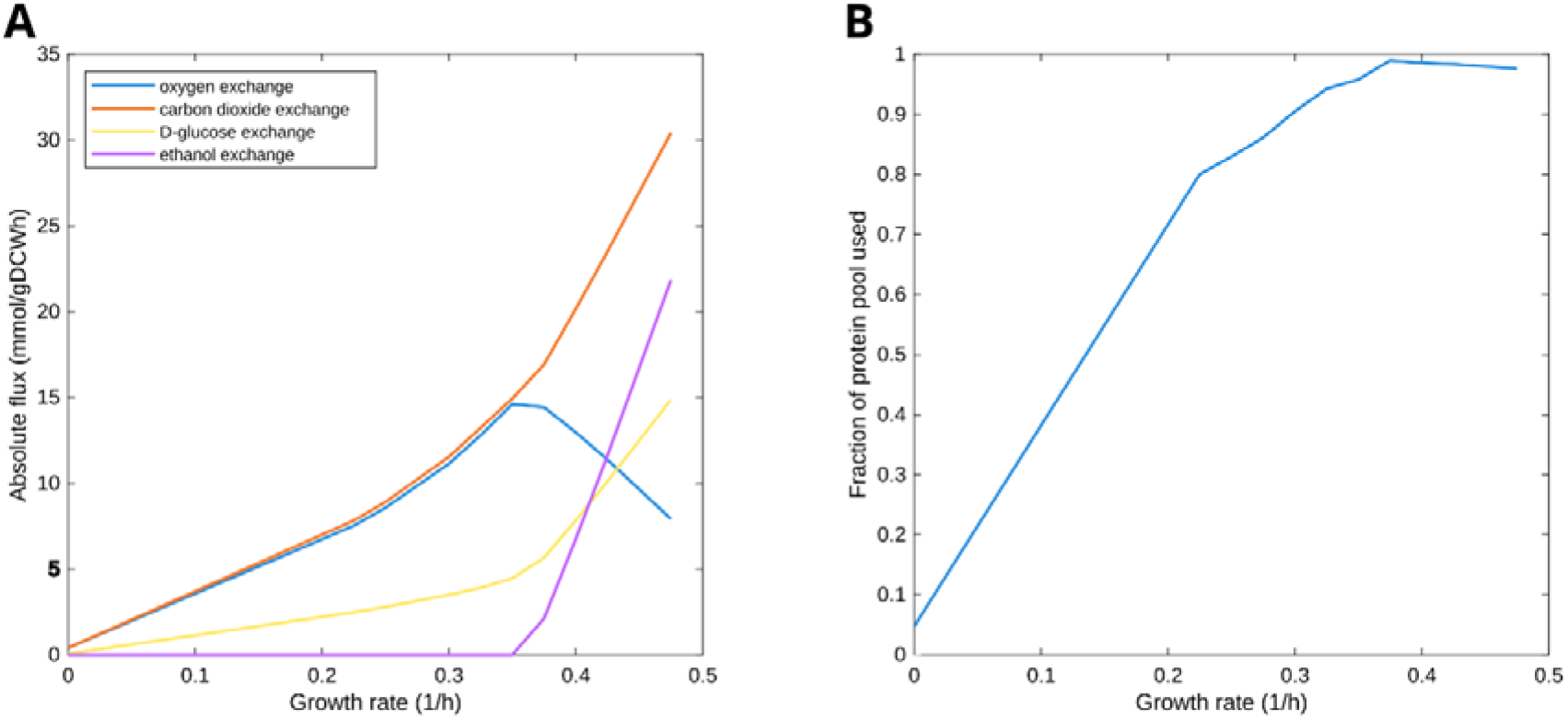
Simulation of chemostats at increasing dilution rates. (A) Absolute flux for oxygen and glucose uptake, and CO2 and ethanol production. (B) Fraction of protein resource use as a function of dilution rate.

These results correlate with those reported on overflow metabolism, where different sets of enzymes are activated and repressed in this metabolic shift, employing enzymes with higher catalytic efficiency to deal with fermentative metabolism. Nevertheless, the observed oxygen uptake at high dilution rates indicates that *K. marxianus*, contrary to *S. cerevisiae* (Crabtree-positive yeast), exhibits a respiro-fermentative type of overflow metabolism, where respiratory and fermentative pathways occur simultaneously at high growth rates. This is agreement with high-sugar batch cultivations of *K. marxianus* conducted in Erlenmeyer flasks, where the cultivations under hypoxia displayed higher lactose consumption and ethanol production than anoxic cultivations, highlighting that lower but still present oxygen content favours the ethanol production from different feedstocks [13,52–55].

### 3.2 Gene co-expression networks reveals that the protein quality control system displays a key role in response to ethanol stress

To investigate the mechanisms affected in the response to ethanol stress in *K. marxianus*, we inferred gene co-expression networks to identify gene modules related to physiological changes. First, we collected transcriptomic data from the literature, using data generated from yeast cultivated under unstressed conditions and exposed to ethanol (stress condition), from two different studies [39,40]. The data from each study were independently used for inferring the GCNs.

In the study conducted by Diniz et al. [39], triplicate samples were collected at three different time intervals: zero hours after 6% (v/v) ethanol exposure (0h, control), one hour after exposure (1h), and four hours after exposure (4h). With the normalized gene count matrix, we first determined the β power value that best satisfied a scale-free topology without drastically reducing average connectivity (Figure S1). The optimal network presented 33 modules (Figure 3). We then performed a Gene Ontology (GO) enrichment analysis for modules positively correlated with the groups. For the 0h condition, the most correlated module contained genes associated with ribosome biogenesis activity (Figure S2). For the 1h condition, the modules featured genes related to glycosylphosphatidylinositol (GPI) anchoring to proteins, nucleolus activity, and ribosomal RNA (rRNA) processing (Figure S3). For the 4h condition, the modules contained genes related to ribosome structure and protein biosynthesis activity (Figure S4).

**Figure 3.**
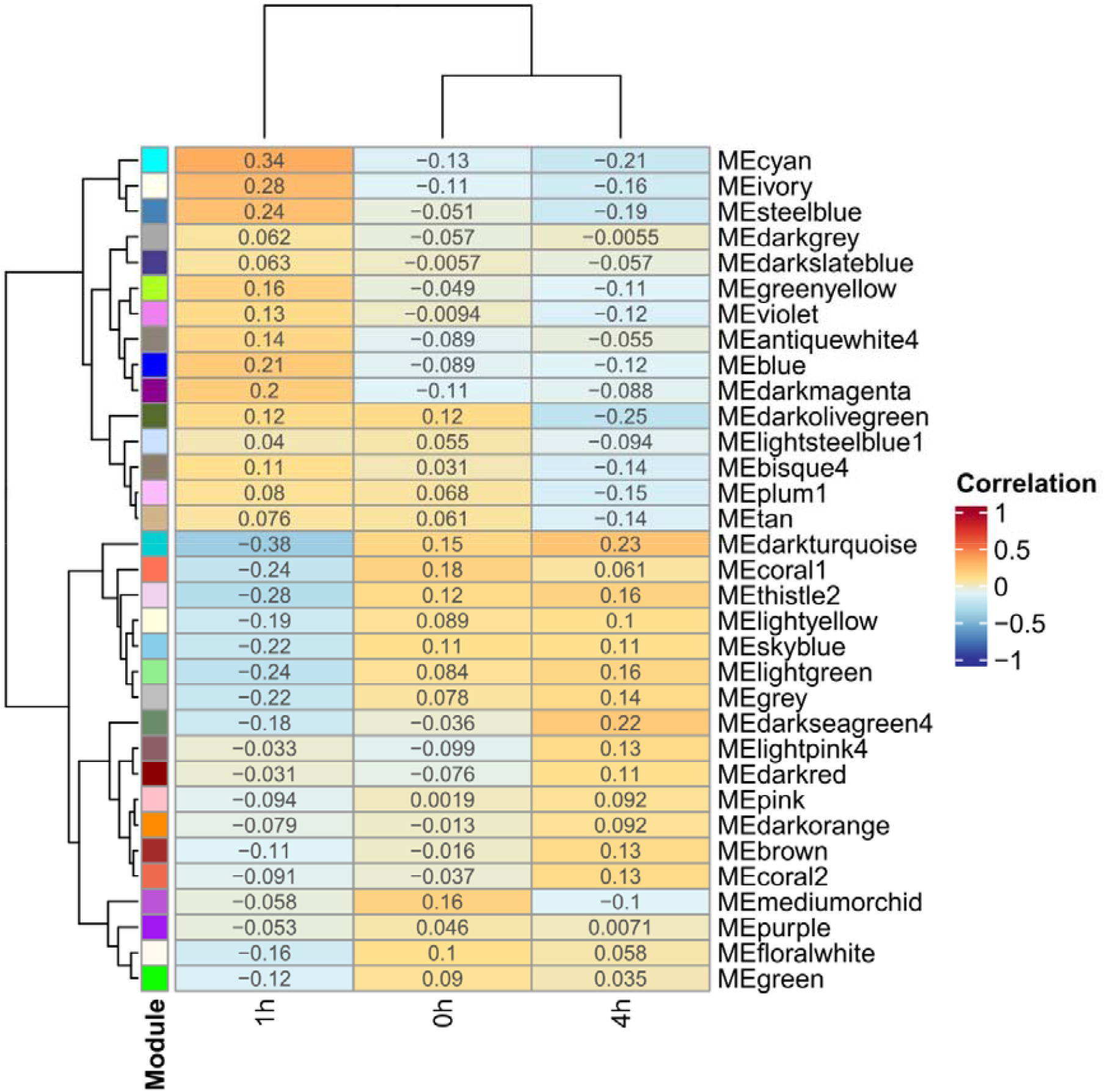
Modules detected and their correlation with the phenotype analysed for the data from Diniz et al. [39].

In the study of Mo et al. [40], laboratory adaptive evolution (ALE) was performed in *K. marxianus* exposed to 6% ethanol (v/v) for 450 generations in order to select evolved strains with improved tolerance to the ethanol stress. For evaluation of the evolved strain, triplicate samples were collected from the following cultures: 0% ethanol (0%-KM, control), 4% ethanol (4%-KM), and 6% ethanol (6%-KM), using the wild-type strain; and 0% ethanol (0%-100d), 4% ethanol (4%-100d), and 6% ethanol (6%-100d) using the evolved strain with higher ethanol tolerance. For inference of the GCNs, we used the count matrix to determine the β power value (Figure S5). The optimal network presented 27 modules (Figure 4). We analysed GO enrichment for modules positively correlated with the groups. For the 0%-KM condition, there were no modules with enriched functions. For 4%-KM, the modules featured genes involved in the citric acid cycle (CAC) and inner membrane of the ribosome (Figure S6). In the 6%-KM condition, the modules contained genes involved in protein transport, post-transcriptional and post-translational modifications, and ubiquitination (Figure S7). The 0%-100d condition also had a module containing genes involved in these same activities (Figure S7A). In the 4%-100d condition, the modules contained genes involved in proteasome activities and protein ubiquitination, microtubules, and kinase regulators (Figure S8). Finally, in the 6%-100d condition, the modules contained genes involved in ribosomal structure and function, protein biosynthesis, and ribosomes (Figure S9). Together, these results indicate that there is a remodelling of the gene expression machinery during ethanol stress, as the modules positively correlated with this condition were mostly enriched with genes related to transcription, translation, and regulation activities. Additionally, in the mutant strains of Mo et al. [40], we observed the presence of genes related to the proteasome, suggesting that protein degradation and recycling are affected during ethanol stress. Since ethanol causes protein denaturation, this could explain why genes related to the gene expression machinery were enriched by certain GO terms, indicating that the protein quality control system is a pivotal mechanism in response to the ethanol stress in *K. marxianus*.

**Figure 4.**
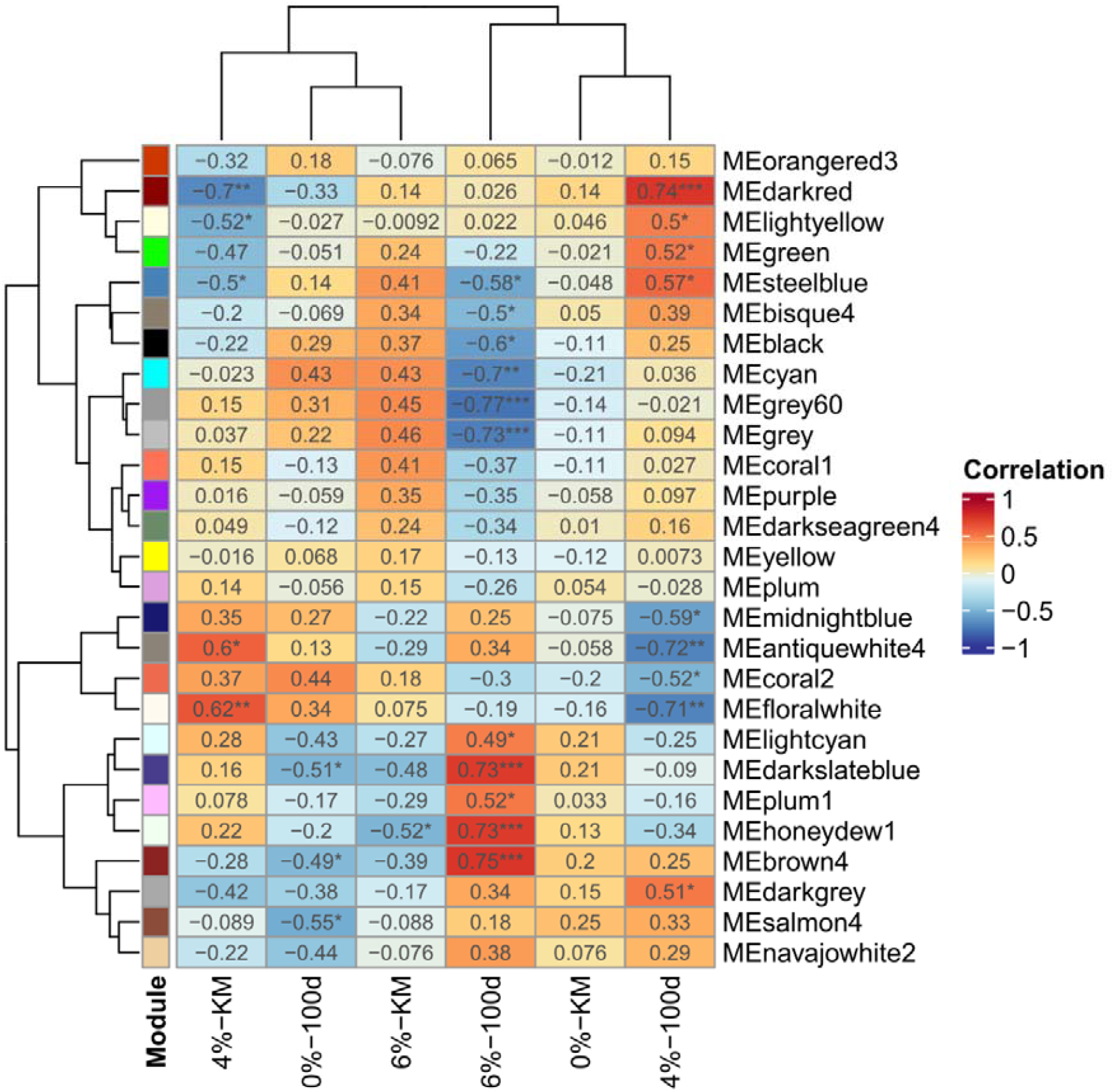
Modules detected and their correlation with the phenotype analysed for the data from Mo et al. [40].

### 3.3 Metabolic impact of transcription factor knockouts

Using the inferred GRNs, we integrated the networks with the iSM996 GEM using PROM to predict how the knockout of each regulator individually impacts metabolism, investigating changes in specific growth rates and flux distributions from the different samples of Diniz et al. [39] and Mo et al. [40].

For the Diniz et al. [39] samples, we observed a total of 13 regulators whose knockout led to a decrease in specific growth rate (Table S1). Among these, notable ones include: the transcriptional activator *HAP3* in the 4h condition, which regulates genes involved in the electron transport chain in the mitochondria; and the transcriptional activator *MET4*, involved in regulating genes for the biosynthesis of sulphur-containing amino acids (methionine, cysteine, and cystine). For the Mo et al. [40] samples, the deletion of all transcription factors resulted in an impact on the specific growth rate (Table S2); notably, the knockout of the regulator *SNF2* led to the lowest specific growth rate among the regulators. This regulator is part of a family of helicase proteins that play a role in chromatin remodelling and is involved in molecular responses to amino acid depletion, response to double-strand DNA breaks, and responses to glucose depletion, all triggered during ethanol stress [14].

### 3.4 Condition-specific models reveal flux redistributions and enzyme activities

To investigate how changes in gene expression affect metabolism, we integrated transcriptomic data with the iSM966 model using RIPTiDe, which restricts the flux of reactions controlled by genes present in the count matrix. We generated condition-specific models for each cultivation condition, from both the Diniz et al. [39] and Mo et al. [40] data. With these condition-specific models, we predicted the corresponding flux distribution, which we then used to constrain the model with enzymatic restriction, eciSM966. With a new round of predictions, we were able to predict the specific growth rate and obtain the flux distribution that approximates to the organism’s physiological reality given the measured gene expression.

The condition-specific models obtained from the Diniz et al. [39] and Mo et al. [40] samples reveal similar phenotypes when compared with the eciSM966 model. Focusing on samples with higher ethanol exposure (6h, 6-KM, 6-100d), we observed that in all three situations, the specific growth rate is low (μ ≤ 0.07). Regarding the flux distribution, metabolic flux was predicted for several reactions related to ethanol metabolism, involving enzymes such as acetaldehyde:NAD+ oxidoreductase and ethanol:NAD+ oxidoreductase. In the 6h condition specifically, we predicted flux for branched-chain amino acid and amino acid production, while in the 6-KM and 6-100d conditions, flux was predicted for ergosterol and membrane lipid biosynthesis. This result is consistent with the results previously described experimentally in the works of Diniz et al. [39] and Mo et al. [40]. Further, the results from the 6-KM and 6-100d conditions also correlate to the ALE experiments of Silveira et al. [56], which have also detected increases in ergosterol and membrane lipid contents in the evolved strains. These evolved strains have also shown an increase in valine (a branched-chain amino acid) and metabolites involved in central pathways, such as isocitric acid, citric acid and cis-aconitic acid, for which the 6h condition-specific model could successfully predict.

## 4 Conclusion

Here we predicted the occurrence of the respiro-fermentative metabolism *in K. marxianus* by using enzyme-constrained metabolic models. Moreover, we obtained new insights about the genes, enzymes, and metabolites involved in the response to ethanol stress in this yeast, integrating transcriptomic data with co-expression network modelling approaches, gene regulatory networks, and metabolic networks. By obtaining results at the resolution of metabolic flux, we identified how metabolism is remodelled to ensure cell survival and the impacts on cell growth. Thus, these results indicate target genes for the implementation of strategies to increase ethanol tolerance, paving the way for further studies of metabolic engineering.

## CRediT authorship contribution statement

### Mauricio Alexander de Moura Ferreira

Conceptualization, Methodology, Software, Formal Analysis, Investigation, Writing - Original Draft, Writing - Review & Editing.

### Wendel Silveira

Conceptualization, Writing - Original Draft, Writing - Review & Editing, Supervision, Project administration, Funding acquisition.

## Conflict of interest statement

The authors declare no conflicts of interest.

### Acknowledgments

This study was financed in part by the Coordenação de Aperfeiçoamento de Pessoal de Nível Superior – Brasil (CAPES) – Finance Code 001 and Conselho Nacional de Desenvolvimento Científico e Tecnológico (CNPq) – Process 312390/2020-3. We thank Eduardo Almeida for his discussions and critical comments on this study.

## Data availability

The code and data used are publicly available in the GitHub repository: https://github.com/LabFisUFV/KmarxianusEthanol. A static version of the repository is available at Zenodo: https://zenodo.org/doi/10.5281/zenodo.11501521.

## Supplementary figures

**Figure S1.**
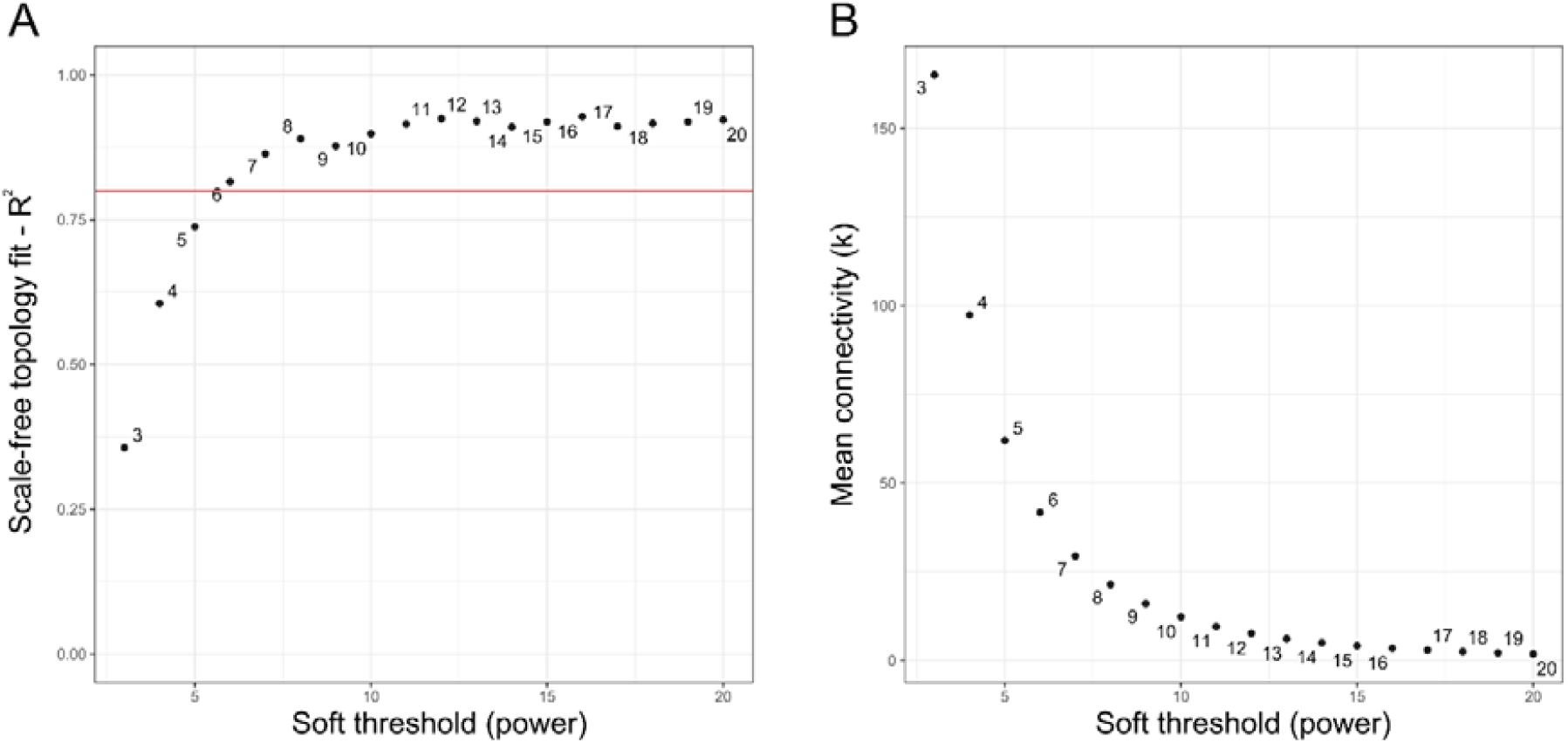
Relationship between scale-free topology and average connectivity for the data from Diniz et al. [39]. The threshold value of 6 was chosen for the subsequent analyses (A) Scale independence. (B) Average connectivity.

**Figure S2.**
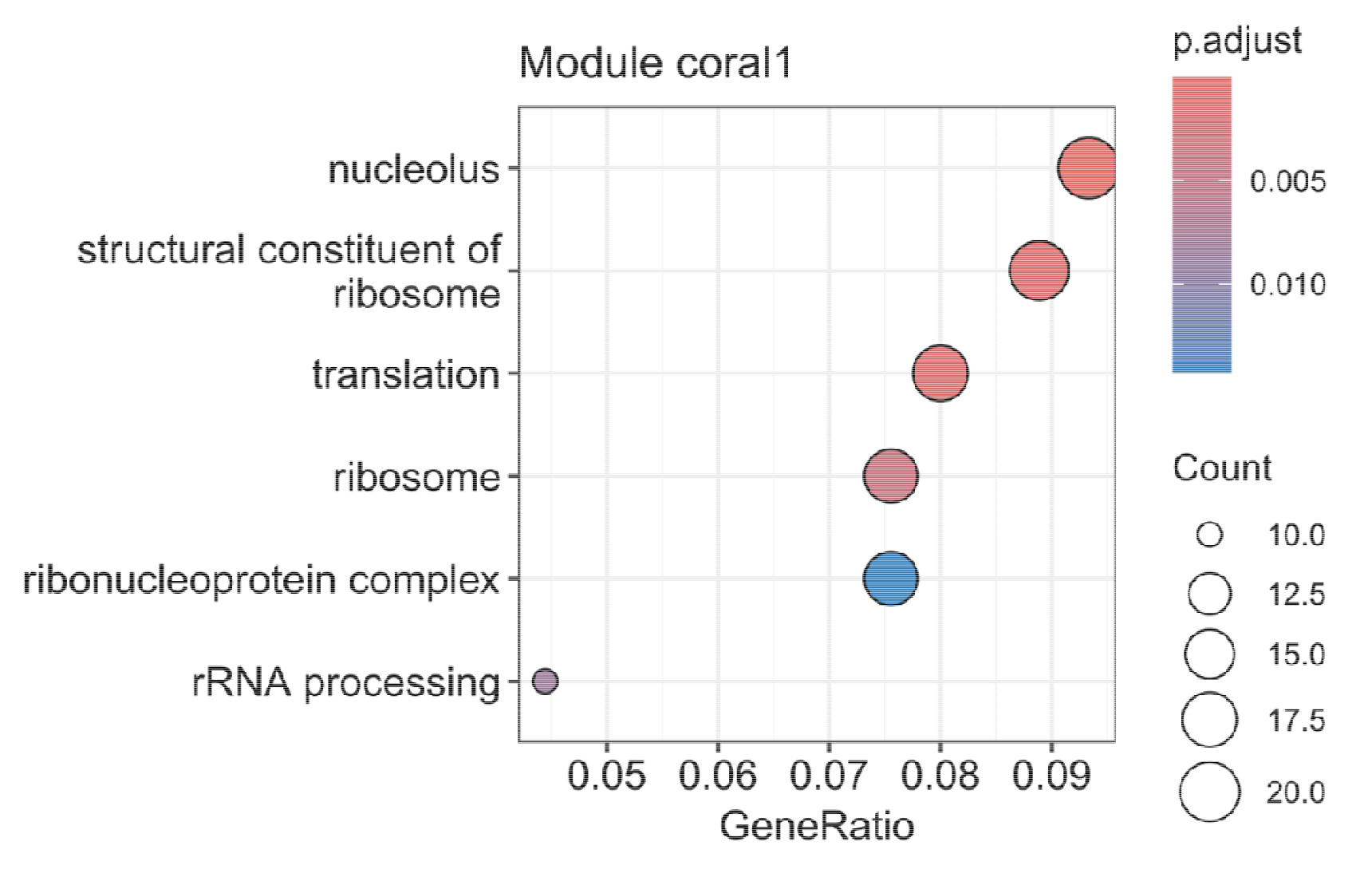
GO enrichment for the coral1 module, the most correlated module in the 0h condition of Diniz et al. [39].

**Figure S3.**
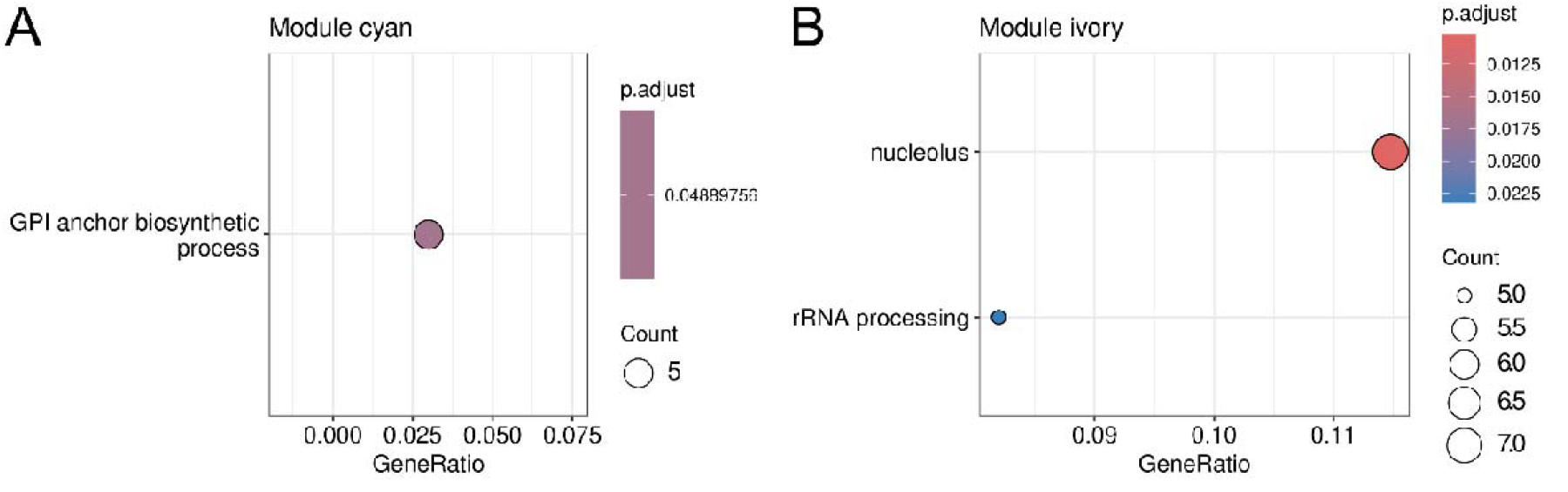
GO enrichment for the cyan (A) and ivory (B) modules, modules most correlated to the 1h condition of Diniz et al. [39].

**Figure S4.**
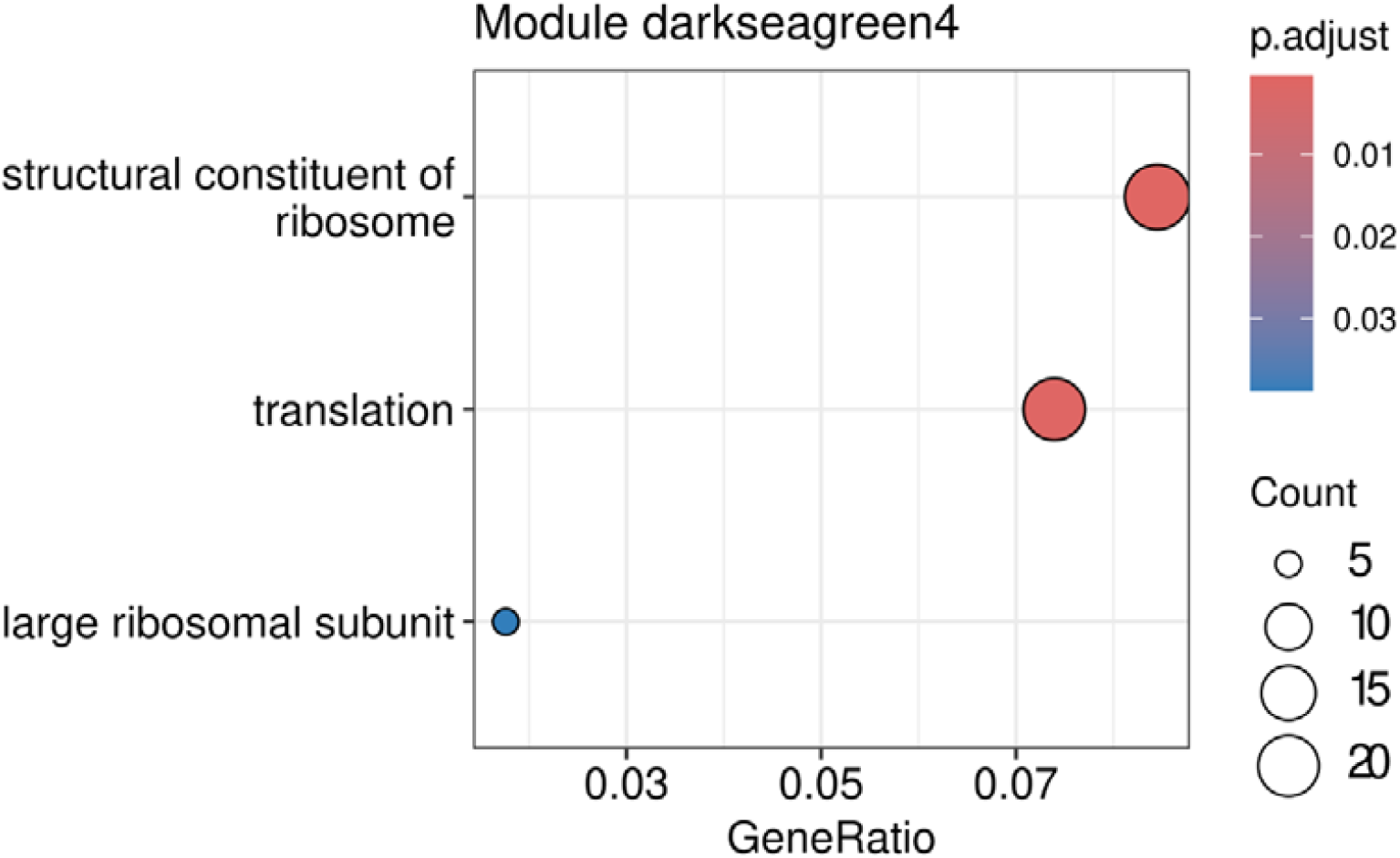
GO enrichment for the darkseagreen4 module, the module most correlated to the 4h condition of Diniz et al. [39].

**Figure S5.**
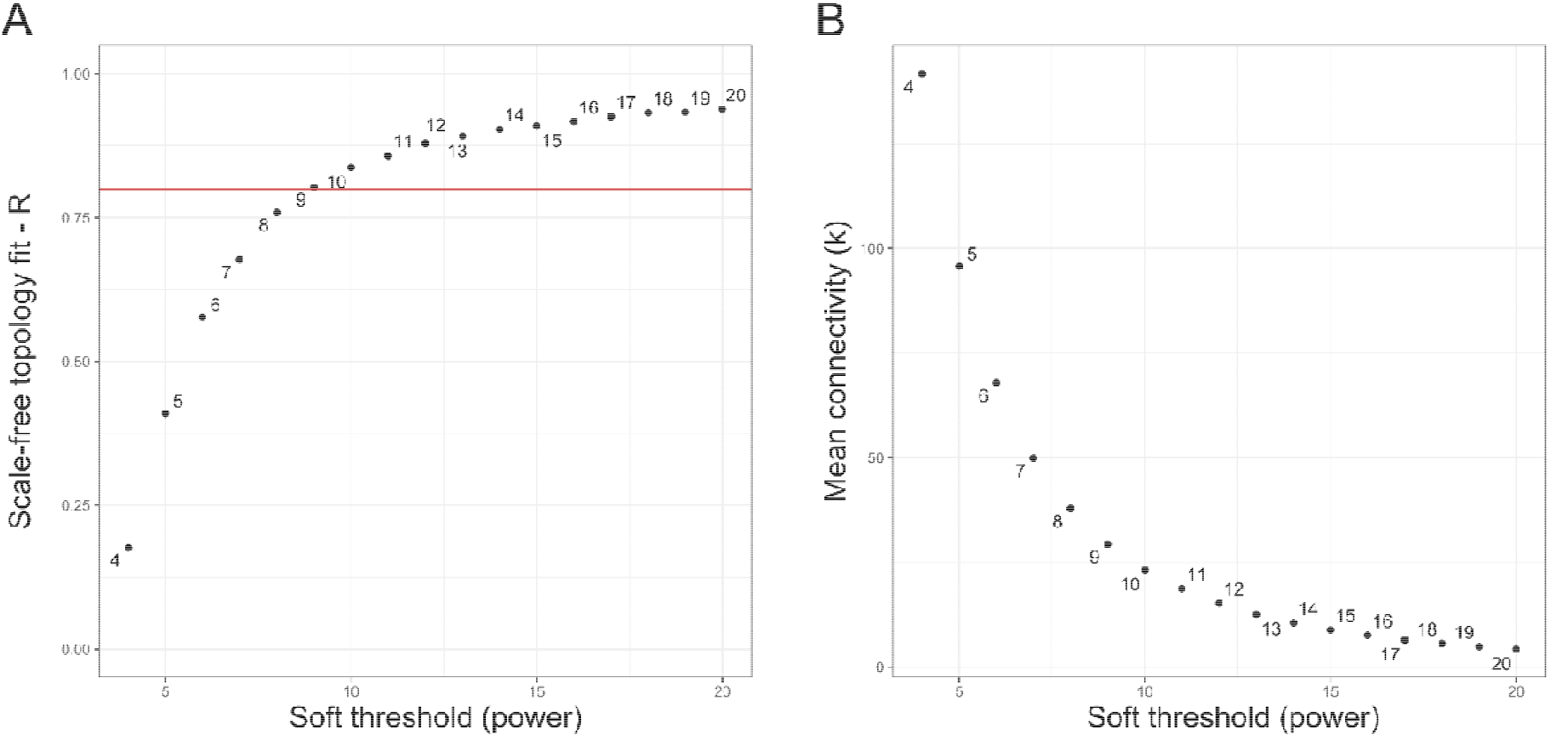
Relationship between scale-free topology and average connectivity for the data from Mo et al. [40]. The threshold value of 9 was chosen for the subsequent analyses (A) Scale independence. (B) Average connectivity.

**Figure S6.**
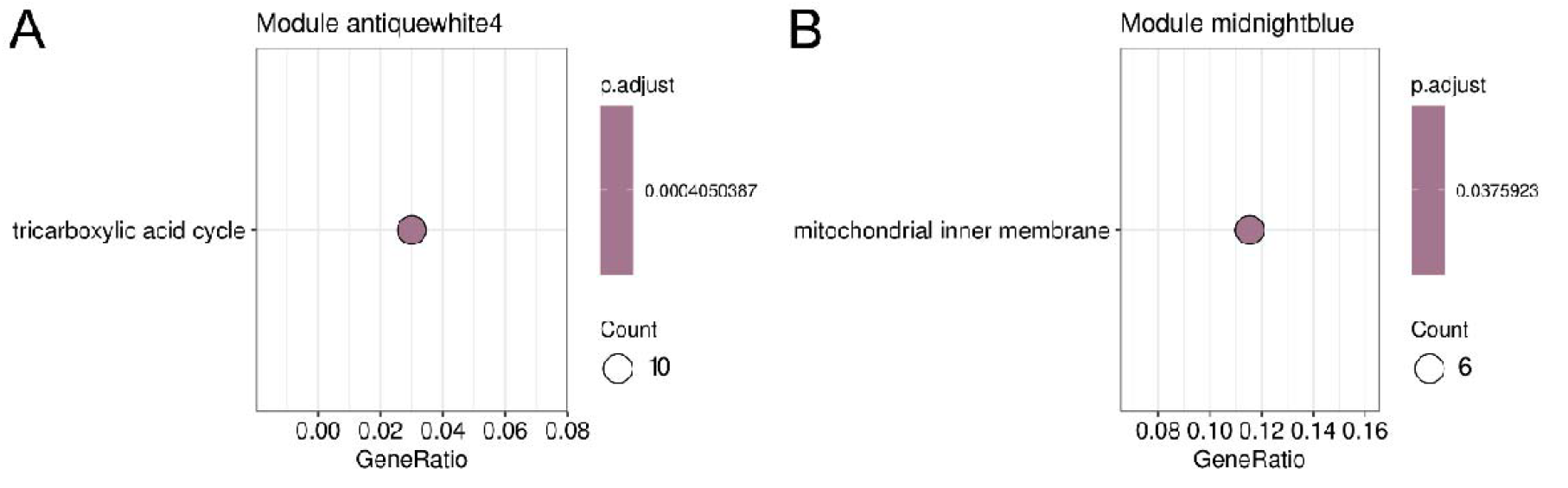
GO enrichment for the antiquewhite4 and midnightblue modules, the most correlated modules in the 4%-KM condition of Mo et al. [40].

**Figure S7.**
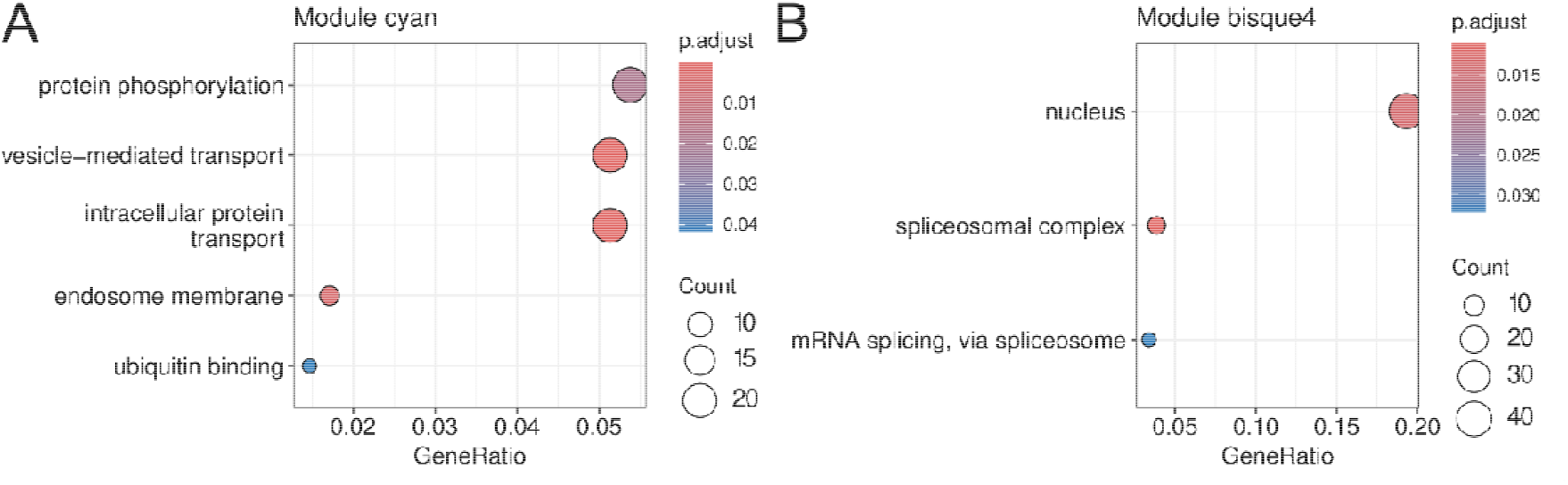
GO enrichment for the cyan (A) and bisque4 (B) modules, the most correlated modules in the 6%-KM (A and B) and 0%-100d (A) condition from Mo et al. [40].

**Figure S8.**
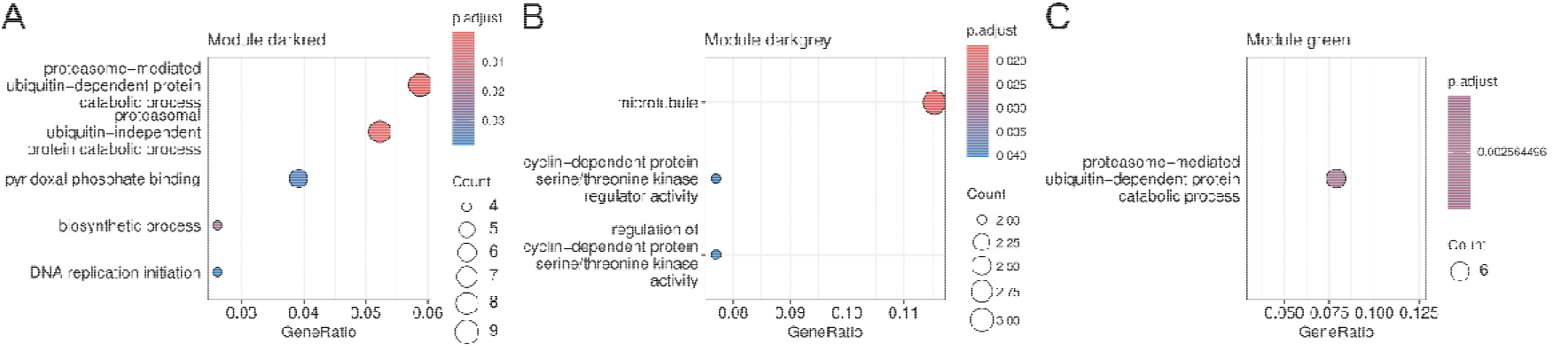
GO enrichment for the darkred (A), darkgrey (B) and green (C) modules, the most correlated modules in the 4%-100d condition of Mo et al. [40].

**Figure S9.**
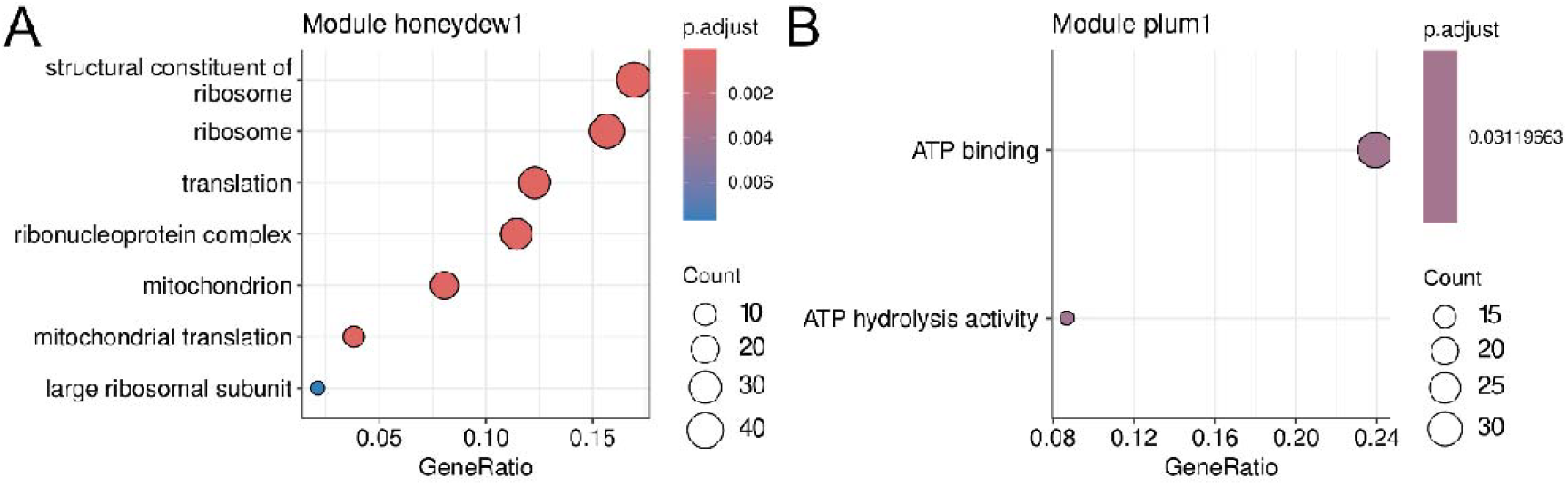
GO enrichment for honeydew1 (A) and plum1 (B), the most correlated modules in the 6%-100d condition from Mo et al. [40].

## References

[1] S.I. Mussatto, G. Dragone, P.M.R. Guimarães, J.P.A. Silva, L.M. Carneiro, I.C. Roberto, A. Vicente, L. Domingues, J.A. Teixeira, Technological trends, global market, and challenges of bio-ethanol production, Biotechnology Advances 28 (2010) 817–830. 10.1016/j.biotechadv.2010.07.001.

[2] M. Unglert, D. Bockey, C. Bofinger, B. Buchholz, G. Fisch, R. Luther, M. Müller, K. Schaper, J. Schmitt, O. Schröder, U. Schümann, H. Tschöke, E. Remmele, R. Wicht, M. Winkler, J. Krahl, Action areas and the need for research in biofuels, Fuel 268 (2020) 117227. 10.1016/j.fuel.2020.117227.

[3] M. Valdivia, J.L. Galan, J. Laffarga, J.L. Ramos, Biofuels 2020: Biorefineries based on lignocellulosic materials, Microbial Biotechnology 9 (2016) 585–594. 10.1111/1751-7915.12387.

[4] RFA, 2020 RFA’s Ethanol Industry Outlook, 2020.

[5] K. Robak, M. Balcerek, Review of second generation bioethanol production from residual biomass, Food Technology and Biotechnology 56 (2018) 174–187. 10.17113/ftb.56.02.18.5428.

[6] I.E.A. Iea, G. Cooper, J. McCaherty, E. Huschitt, R. Schwarck, C. Wilson, 2021 Ethanol Industry Outlook, Renewable Fuels Association (2021) 1–40.

[7] S. Moreno-Paz, J. Schmitz, V.A.P. Martins dos Santos, M. Suarez-Diez, Enzyme-constrained models predict the dynamics of Saccharomyces cerevisiae growth in continuous, batch and fed-batch bioreactors, Microbial Biotechnology 15 (2022) 1434–1445. 10.1111/1751-7915.13995.

[8] E. de Alteriis, F. Cartenì, P. Parascandola, J. Serpa, S. Mazzoleni, Revisiting the Crabtree/Warburg effect in a dynamic perspective: a fitness advantage against sugar-induced cell death, Cell Cycle 17 (2018) 688–701. 10.1080/15384101.2018.1442622.

[9] Y.J. Liu, B. Li, Y. Feng, Q. Cui, Consolidated bio-saccharification: Leading lignocellulose bioconversion into the real world, Biotechnology Advances 40 (2020) 107535. 10.1016/j.biotechadv.2020.107535.

[10] M.M. Lane, J.P. Morrissey, Kluyveromyces marxianus: A yeast emerging from its sister’s shadow, Fungal Biology Reviews 24 (2010) 17–26. 10.1016/j.fbr.2010.01.001.

[11] M. Arellano-Plaza, R. Noriega-Cisneros, M. Clemente-Guerrero, J.C. González-Hernández, P.D. Robles-Herrera, S. Manzo-Ávalos, A. Saavedra-Molina, A. Gschaedler-Mathis, Fermentative capacity of Kluyveromyces marxianus and Saccharomyces cerevisiae after oxidative stress, Journal of the Institute of Brewing 123 (2017) 519–526. 10.1002/jib.451.

[12] D.A. Costa, C.J.A. de Souza, P.A. Costa, M.Q.R.B. Rodrigues, A.F. dos Santos, M.R. Lopes, H.L.A. Genier, W.B. Silveira, L.G. Fietto, Physiological characterization of thermotolerant yeast for cellulosic ethanol production., Applied Microbiology and Biotechnology 98 (2014) 3829–40. 10.1007/s00253-014-5580-3.

[13] W.B. Silveira, F.J.V. Passos, H.C. Mantovani, F.M.L. Passos, Ethanol production from cheese whey permeate by Kluyveromyces marxianus UFV-3: A flux analysis of oxido-reductive metabolism as a function of lactose concentration and oxygen levels, Enzyme and Microbial Technology 36 (2005) 930–936. 10.1016/j.enzmictec.2005.01.018.

[14] M.A. de Moura Ferreira, F.A. da Silveira, W.B. da Silveira, Ethanol stress responses in Kluyveromyces marxianus: current knowledge and perspectives, Appl Microbiol Biotechnol 106 (2022) 1341–1353. 10.1007/s00253-022-11799-0.

[15] X. Fu, P. Li, L. Zhang, S. Li, Understanding the stress responses of Kluyveromyces marxianus after an arrest during high-temperature ethanol fermentation based on integration of RNA-Seq and metabolite data, Applied Microbiology and Biotechnology 103 (2019) 2715–2729. 10.1007/s00253-019-09637-x.

[16] F. Vriesekoop, C. Haass, N.B. Pamment, The role of acetaldehyde and glycerol in the adaptation to ethanol stress of Saccharomyces cerevisiae and other yeasts, FEMS Yeast Research 9 (2009) 365–371. 10.1111/j.1567-1364.2009.00492.x.

[17] J.D. Orth, I. Thiele, B.Ø. Palsson, What is flux balance analysis?, Nature Biotechnology 28 (2010) 245–248. 10.1038/nbt.1614.

[18] R. Adadi, B. Volkmer, R. Milo, M. Heinemann, T. Shlomi, Prediction of Microbial Growth Rate versus Biomass Yield by a Metabolic Network with Kinetic Parameters, PLOS Computational Biology 8 (2012) e1002575. 10.1371/JOURNAL.PCBI.1002575.

[19] B.J. Sánchez, C. Zhang, A. Nilsson, P.-J. Lahtvee, E.J. Kerkhoven, J. Nielsen, Improving the phenotype predictions of a yeast genome-scale metabolic model by incorporating enzymatic constraints, Molecular Systems Biology 13 (2017) 935. 10.15252/MSB.20167411.

[20] W. Guo, X. Feng, OM-FBA: Integrate Transcriptomics Data with Flux Balance Analysis to Decipher the Cell Metabolism, PLOS ONE 11 (2016) e0154188. 10.1371/journal.pone.0154188.

[21] C. Ruprecht, N. Vaid, S. Proost, S. Persson, M. Mutwil, Beyond Genomics: Studying Evolution with Gene Coexpression Networks, Trends in Plant Science 22 (2017) 298–307. 10.1016/j.tplants.2016.12.011.

[22] M. Zhao, W. He, J. Tang, Q. Zou, F. Guo, A comprehensive overview and critical evaluation of gene regulatory network inference technologies, Briefings in Bioinformatics 00 (2021) 1–15. 10.1093/bib/bbab009.

[23] S. Marcišauskas, B. Ji, J. Nielsen, Reconstruction and analysis of a Kluyveromyces marxianus genome-scale metabolic model, BMC Bioinformatics 20 (2019) 1–9. 10.1186/s12859-019-3134-5.

[24] Y. Chen, J. Gustafsson, A. Tafur Rangel, M. Anton, I. Domenzain, C. Kittikunapong, F. Li, L. Yuan, J. Nielsen, E.J. Kerkhoven, Reconstruction, simulation and analysis of enzyme-constrained metabolic models using GECKO Toolbox 3.0, Nat Protoc 19 (2024) 629–667. 10.1038/s41596-023-00931-7.

[25] T.U. Consortium, UniProt: a worldwide hub of protein knowledge, Nucleic Acids Research 47 (2018) D506–D515. 10.1093/nar/gky1049.

[26] M. Kanehisa, S. Goto, KEGG: Kyoto Encyclopedia of Genes and Genomes, Nucleic Acids Research 28 (2000) 27–30. 10.1093/nar/28.1.27.

[27] A. Chang, L. Jeske, S. Ulbrich, J. Hofmann, J. Koblitz, I. Schomburg, M. Neumann-Schaal, D. Jahn, D. Schomburg, BRENDA, the ELIXIR core data resource in 2021: new developments and updates, Nucleic Acids Research 49 (2021) D498–D508. 10.1093/NAR/GKAA1025.

[28] F. Li, L. Yuan, H. Lu, G. Li, Y. Chen, M.K.M. Engqvist, E.J. Kerkhoven, J. Nielsen, Deep learning-based kcat prediction enables improved enzyme-constrained model reconstruction, Nature Catalysis (2022) 1–11. 10.1038/s41929-022-00798-z.

[29] S. Kim, J. Chen, T. Cheng, A. Gindulyte, J. He, S. He, Q. Li, B.A. Shoemaker, P.A. Thiessen, B. Yu, L. Zaslavsky, J. Zhang, E.E. Bolton, PubChem 2023 update, Nucleic Acids Research 51 (2023) D1373–D1380. 10.1093/nar/gkac956.

[30] G.G. Fonseca, N.M.B. de Carvalho, A.K. Gombert, Growth of the yeast Kluyveromyces marxianus CBS 6556 on different sugar combinations as sole carbon and energy source, Appl Microbiol Biotechnol 97 (2013) 5055–5067. 10.1007/s00253-013-4748-6.

[31] L.H. Bellaver, N.M.B. de Carvalho, J. Abrahão-Neto, A.K. Gombert, Ethanol formation and enzyme activities around glucose-6-phosphate in Kluyveromyces marxianus CBS 6556 exposed to glucose or lactose excess, FEMS Yeast Res 4 (2004) 691–698. 10.1016/j.femsyr.2004.01.004.

[32] G.G. Fonseca, A.K. Gombert, E. Heinzle, C. Wittmann, Physiology of the yeast Kluyveromyces marxianus during batch and chemostat cultures with glucose as the sole carbon source, FEMS Yeast Research 7 (2007) 422–435. 10.1111/j.1567-1364.2006.00192.x.

[33] E. Postma, P.J. Van den Broek, Continuous-culture study of the regulation of glucose and fructose transport in Kluyveromyces marxianus CBS 6556, Journal of Bacteriology 172 (1990) 2871–2876. 10.1128/jb.172.6.2871-2876.1990.

[34] A. Pentjuss, E. Stalidzans, J. Liepins, A. Kokina, J. Martynova, P. Zikmanis, I. Mozga, R. Scherbaka, H. Hartman, M.G. Poolman, D.A. Fell, A. Vigants, Model-based biotechnological potential analysis of Kluyveromyces marxianus central metabolism, Journal of Industrial Microbiology and Biotechnology 44 (2017) 1177–1190. 10.1007/s10295-017-1946-8.

[35] L.G.S. Longhi, D.J. Luvizetto, L.S. Ferreira, R. Rech, M.A.Z. Ayub, A.R. Secchi, A growth kinetic model of Kluyveromyces marxianus cultures on cheese whey as substrate, J IND MICROBIOL BIOTECHNOL 31 (2004) 35–40. 10.1007/s10295-004-0110-4.

[36] M. Hensing, H. Vrouwenvelder, C. Hellinga, R. Baartmans, H. van Dijken, Production of extracellular inulinase in high-cell-density fed-batch cultures of Kluyveromyces marxianus, Appl Microbiol Biotechnol 42 (1994) 516–521. 10.1007/BF00173914.

[37] L. Heirendt, S. Arreckx, T. Pfau, S.N. Mendoza, A. Richelle, A. Heinken, H.S. Haraldsdóttir, J. Wachowiak, S.M. Keating, V. Vlasov, S. Magnusdóttir, C.Y. Ng, G. Preciat, A. Žagare, S.H.J. Chan, M.K. Aurich, C.M. Clancy, J. Modamio, J.T. Sauls, A. Noronha, A. Bordbar, B. Cousins, D.C. El Assal, L.V. Valcarcel, I. Apaolaza, S. Ghaderi, M. Ahookhosh, M. Ben Guebila, A. Kostromins, N. Sompairac, H.M. Le, D. Ma, Y. Sun, L. Wang, J.T. Yurkovich, M.A.P. Oliveira, P.T. Vuong, L.P. El Assal, I. Kuperstein, A. Zinovyev, H.S. Hinton, W.A. Bryant, F.J. Aragón Artacho, F.J. Planes, E. Stalidzans, A. Maass, S. Vempala, M. Hucka, M.A. Saunders, C.D. Maranas, N.E. Lewis, T. Sauter, B. Palsson, I. Thiele, R.M.T. Fleming, Creation and analysis of biochemical constraint-based models using the COBRA Toolbox v.3.0, Nature Protocols 14 (2019) 639–702. 10.1038/s41596-018-0098-2.

[38] H. Wang, S. Marcišauskas, B.J. Sánchez, I. Domenzain, D. Hermansson, R. Agren, J. Nielsen, E.J. Kerkhoven, RAVEN 2.0: A versatile toolbox for metabolic network reconstruction and a case study on Streptomyces coelicolor, PLOS Computational Biology 14 (2018) e1006541. 10.1371/journal.pcbi.1006541.

[39] R.H.S. Diniz, J.C. Villada, M.C.T. Alvim, P.M.P. Vidigal, N.M. Vieira, M. Lamas-Maceiras, M.E. Cerdán, M.I. González-Siso, P.J. Lahtvee, W.B. da Silveira, Transcriptome analysis of the thermotolerant yeast Kluyveromyces marxianus CCT 7735 under ethanol stress, Applied Microbiology and Biotechnology 101 (2017) 6969–6980. 10.1007/s00253-017-8432-0.

[40] W. Mo, M. Wang, R. Zhan, Y. Yu, Y. He, H. Lu, Kluyveromyces marxianus developing ethanol tolerance during adaptive evolution with significant improvements of multiple pathways, Biotechnology for Biofuels 12 (2019) 1–15. 10.1186/s13068-019-1393-z.

[41] B. Langmead, S.L. Salzberg, Fast gapped-read alignment with Bowtie 2, Nat Methods 9 (2012) 357–359. 10.1038/nmeth.1923.

[42] Y. Liao, G.K. Smyth, W. Shi, featureCounts: an efficient general purpose program for assigning sequence reads to genomic features, Bioinformatics 30 (2014) 923–930. 10.1093/bioinformatics/btt656.

[43] M.I. Love, W. Huber, S. Anders, Moderated estimation of fold change and dispersion for RNA-seq data with DESeq2, Genome Biol 15 (2014) 1–21. 10.1186/s13059-014-0550-8.

[44] F. Almeida-Silva, T.M. Venancio, BioNERO: an all-in-one R/Bioconductor package for comprehensive and easy biological network reconstruction, Funct Integr Genomics 22 (2022) 131–136. 10.1007/s10142-021-00821-9.

[45] R.C. Gentleman, V.J. Carey, D.M. Bates, B. Bolstad, M. Dettling, S. Dudoit, B. Ellis, L. Gautier, Y. Ge, J. Gentry, K. Hornik, T. Hothorn, W. Huber, S. Iacus, R. Irizarry, F. Leisch, C. Li, M. Maechler, A.J. Rossini, G. Sawitzki, C. Smith, G. Smyth, L. Tierney, J.Y. Yang, J. Zhang, Bioconductor: open software development for computational biology and bioinformatics, Genome Biol 5 (2004) 1–16. 10.1186/gb-2004-5-10-r80.

[46] S. Carbon, E. Douglass, B.M. Good, D.R. Unni, N.L. Harris, C.J. Mungall, S. Basu, R.L. Chisholm, R.J. Dodson, E. Hartline, P. Fey, P.D. Thomas, L.P. Albou, D. Ebert, M.J. Kesling, H. Mi, A. Muruganujan, X. Huang, T. Mushayahama, S.A. LaBonte, D.A. Siegele, G. Antonazzo, H. Attrill, N.H. Brown, P. Garapati, S.J. Marygold, V. Trovisco, G. dos Santos, K. Falls, C. Tabone, P. Zhou, J.L. Goodman, V.B. Strelets, J. Thurmond, P. Garmiri, R. Ishtiaq, M. Rodríguez-López, M.L. Acencio, M. Kuiper, A. Lægreid, C. Logie, R.C. Lovering, B. Kramarz, S.C.C. Saverimuttu, S.M. Pinheiro, H. Gunn, R. Su, K.E. Thurlow, M. Chibucos, M. Giglio, S. Nadendla, J. Munro, R. Jackson, M.J. Duesbury, N. Del-Toro, B.H.M. Meldal, K. Paneerselvam, L. Perfetto, P. Porras, S. Orchard, A. Shrivastava, H.Y. Chang, R.D. Finn, A.L. Mitchell, N.D. Rawlings, L. Richardson, A. Sangrador-Vegas, J.A. Blake, K.R. Christie, M.E. Dolan, H.J. Drabkin, D.P. Hill, L. Ni, D.M. Sitnikov, M.A. Harris, S.G. Oliver, K. Rutherford, V. Wood, J. Hayles, J. Bähler, E.R. Bolton, J.L. de Pons, M.R. Dwinell, G.T. Hayman, M.L. Kaldunski, A.E. Kwitek, S.J.F. Laulederkind, C. Plasterer, M.A. Tutaj, M. Vedi, S.J. Wang, P. D’Eustachio, L. Matthews, J.P. Balhoff, S.A. Aleksander, M.J. Alexander, J.M. Cherry, S.R. Engel, F. Gondwe, K. Karra, S.R. Miyasato, R.S. Nash, M. Simison, M.S. Skrzypek, S. Weng, E.D. Wong, M. Feuermann, P. Gaudet, A. Morgat, E. Bakker, T.Z. Berardini, L. Reiser, S. Subramaniam, E. Huala, C.N. Arighi, A. Auchincloss, K. Axelsen, G. Argoud-Puy, A. Bateman, M.C. Blatter, E. Boutet, E. Bowler, L. Breuza, A. Bridge, R. Britto, H. Bye-A-Jee, C.C. Casas, E. Coudert, P. Denny, A. Es-Treicher, M.L. Famiglietti, G. Georghiou, A.N. Gos, N. Gruaz-Gumowski, E. Hatton-Ellis, C. Hulo, A. Ignatchenko, F. Jungo, K. Laiho, P. Le Mercier, D. Lieberherr, A. Lock, Y. Lussi, A. MacDougall, M. Ma-Grane, M.J. Martin, P. Masson, D.A. Natale, N. Hyka-Nouspikel, S. Orchard, I. Pedruzzi, L. Pourcel, S. Poux, S. Pundir, C. Rivoire, E. Speretta, S. Sundaram, N. Tyagi, K. Warner, R. Zaru, C.H. Wu, A.D. Diehl, J.N. Chan, C. Grove, R.Y.N. Lee, H.M. Muller, D. Raciti, K. van Auken, P.W. Sternberg, M. Berriman, M. Paulini, K. Howe, S. Gao, A. Wright, L. Stein, D.G. Howe, S. Toro, M. Westerfield, P. Jaiswal, L. Cooper, J. Elser, The Gene Ontology resource: enriching a GOld mine, Nucleic Acids Research 49 (2021) D325–D334. 10.1093/NAR/GKAA1113.

[47] P. Jones, D. Binns, H.-Y. Chang, M. Fraser, W. Li, C. McAnulla, H. McWilliam, J. Maslen, A. Mitchell, G. Nuka, S. Pesseat, A.F. Quinn, A. Sangrador-Vegas, M. Scheremetjew, S.-Y. Yong, R. Lopez, S. Hunter, InterProScan 5: genome-scale protein function classification., Bioinformatics (Oxford, England) 30 (2014) 1236–40. 10.1093/bioinformatics/btu031.

[48] M.C. Teixeira, R. Viana, M. Palma, J. Oliveira, M. Galocha, M.N. Mota, D. Couceiro, M.G. Pereira, M. Antunes, I.V. Costa, P. Pais, C. Parada, C. Chaouiya, I. Sá-Correia, P.T. Monteiro, YEASTRACT+: a portal for the exploitation of global transcription regulation and metabolic model data in yeast biotechnology and pathogenesis, Nucleic Acids Research 51 (2023) D785–D791. 10.1093/nar/gkac1041.

[49] S. Chandrasekaran, N.D. Price, Probabilistic integrative modeling of genome-scale metabolic and regulatory networks in Escherichia coli and Mycobacterium tuberculosis, Proceedings of the National Academy of Sciences of the United States of America 107 (2010) 17845–17850. 10.1073/pnas.1005139107.

[50] M.L. Jenior, T.J.M. Jr, B.V. Dougherty, J.A. Papin, Transcriptome-guided parsimonious flux analysis improves predictions with metabolic networks in complex environments, PLOS Computational Biology 16 (2020) e1007099. 10.1371/journal.pcbi.1007099.

[51] Q.K. Beg, A. Vazquez, J. Ernst, M.A. De Menezes, Z. Bar-Joseph, A.L. Barabási, Z.N. Oltvai, Intracellular crowding defines the mode and sequence of substrate uptake by Escherichia coli and constrains its metabolic activity, Proceedings of the National Academy of Sciences of the United States of America 104 (2007) 12663–12668. 10.1073/pnas.0609845104.

[52] R.H.S. Diniz, W.B. Silveira, L.G. Fietto, F.M.L. Passos, The high fermentative metabolism of Kluyveromyces marxianus UFV-3 relies on the increased expression of key lactose metabolic enzymes, Antonie van Leeuwenhoek, International Journal of General and Molecular Microbiology 101 (2012) 541–550. 10.1007/s10482-011-9668-9.

[53] R.H.S. Diniz, M.Q.R.B. Rodrigues, L.G. Fietto, F.M.L. Passos, W.B. Silveira,Optimizing and validating the production of ethanol from cheese whey permeate by Kluyveromyces marxianus UFV-3, Biocatalysis and Agricultural Biotechnology 3 (2014) 111–117. 10.1016/j.bcab.2013.09.002.

[54] P.G. Ferreira, F.A. da Silveira, R.C.V. dos Santos, H.L.A. Genier, R.H.S. Diniz, J.I. Ribeiro, L.G. Fietto, F.M.L. Passos, W.B. da Silveira, Optimizing ethanol production by thermotolerant Kluyveromyces marxianus CCT 7735 in a mixture of sugarcane bagasse and ricotta whey, Food Science and Biotechnology 24 (2015) 1421–1427. 10.1007/s10068-015-0182-0.

[55] L. Signori, S. Passolunghi, L. Ruohonen, D. Porro, P. Branduardi, Effect of oxygenation and temperature on glucose-xylose fermentation in Kluyveromyces marxianus CBS712 strain, Microb Cell Fact 13 (2014) 51. 10.1186/1475-2859-13-51.

[56] F.A. Silveira, D.L. de Oliveira Soares, K.W. Bang, T.R. Balbino, M.A. de Moura Ferreira, R.H.S. Diniz, L.A. de Lima, M.M. Brandão, S.G. Villas-Bôas, W.B. da Silveira, Assessment of ethanol tolerance of Kluyveromyces marxianus CCT 7735 selected by adaptive laboratory evolution, Applied Microbiology and Biotechnology 104 (2020) 7483–7494. 10.1007/s00253-020-10768-9.

